# Impact of repeated cryopreservation on embryo health and implantation potential

**DOI:** 10.1101/2023.10.26.564306

**Authors:** Tong Li, Shan Li, Darren J.X. Chow, Ryan D. Rose, Tiffany C.Y. Tan, Kylie R. Dunning

**Affiliations:** Robinson Research Institute, School of Biomedicine, The University of Adelaide, Adelaide, Australia; Institute for Photonics and Advanced Sensing, The University of Adelaide, Adelaide, Australia; Genea Fertility SA, St. Andrews Hospital, Adelaide, SA, Australia

**Keywords:** Cryopreservation, embryo health, implantation potential, embryo transfer

## Abstract

In IVF clinics, preimplantation genetic testing (PGT) is a common practice that involves a biopsy and cryopreservation of embryos for genetic evaluation. When testing fails – or is required for already cryopreserved embryos – multiple freeze-thaw cycles occur. Though known to impact live birth rates, the exclusive influence of cryopreservation has not been elucidated. Here, we evaluate the effect of repeated cryopreservation on embryo health and implantation potential.

Blastocyst-stage murine embryos were subjected to one, two or three freeze-thaw cycles with fresh embryos serving as a control. Outcomes assessed included post-thaw survival rate, allocation of cells to the inner cell mass (ICM) *vs*. trophectoderm cell lineages, implantation potential and offspring health.

Post-thawing, embryos that were subjected to three freeze-thaw cycles had a significantly lower survival rates compared to embryos that had undergone one cycle (*P*<0.001). Additionally, the number of ICM cells was significantly reduced in embryos subjected to two or three freeze-thaw cycles compared to fresh or single-cycle embryos (*P*<0.001). No statistically significant differences were found for pregnancy rate, number of implantations, viable fetuses or resorption sites between treatment groups. We did however, find a non-significant yet interesting trend: three freeze-thaw cycles were associated with a 20% decrease in viable fetuses and a 20% increase in resorption sites compared to one freeze-thaw cycle group.

These findings demonstrate that repeated cryopreservation adversely affects embryo health and may decrease implantation potential. Consequently, caution is advised regarding the repeated application of cryopreservation in IVF clinics, underscoring the need for further research to optimise cryopreservation protocols.

## Introduction

Infertility is defined as the inability of a couple to achieve a successful pregnancy outcome following 12 months of unprotected intercourse^1^. Nowadays, increasing maternal age, along with the impact of lifestyle factors including obesity and diabetes all contribute to the fact that one in six Australian couples have trouble conceiving naturally^1^. Globally, it is estimated that 48 million couples suffer from infertility^1^. Assisted reproductive technologies (ART) were established as a treatment for infertility ^2–4^.

Cryopreservation is one of a suite of ART that is widely practiced in fertility clinics. This technology allows for the preservation of gametes or embryos at ultra-low temperatures for long-term storage, permitting their future use by patients^5^. This includes patients who wish to delay childbearing, or in situations where their reproductive potential has been compromised by medical pathology^6^. While many studies have showed that vitrification, an ultra-rapid cryopreservation procedure, has comparable pregnancy rates and postnatal outcomes compared to fresh embryo transfers^7–11^, there are contradictory studies that showed detrimental impact on clinical pregnancy rate and live birth rate^12–13^. Additionally, a recent study in an ovine model showed that cryopreserved embryos are prone to apoptosis and have lower cryosurvival rates compared to non-cryopreserved embryos^14^. However, the impact of cryopreservation on the developmental and implantation potential of embryo remains unclear.

In addition to long-term storage of embryos, cryopreservation is also commonly used in conjunction with preimplantation genetic testing for aneuploidy (PGT-A), which is a well-established procedure designed to identify aneuploidy^15–16^. Aneuploidy refers to a cell that contains an unbalanced number of chromosomes, while euploidy refers to when a cell contains the expected number of chromosomes, 46 in humans and 40 in mice. PGT-A involves taking a small biopsy of the trophectoderm cells (placental lineage) at the blastocyst-stage of embryo development. These biopsied cells are sequenced to detect chromosomal abnormalities (aneuploidy) whilst the remaining embryo is vitrified and stored.

Upon receipt of favourable PGT-A results, the embryo is revived and transferred into the uterus of the patient^17^. Importantly, there is an established correlation between a higher proportion of aneuploid cells in the embryo and decreased pregnancy success^18^. By identifying and eliminating the effects of aneuploidy, PGT-A has a positive effect on the pregnancy rates in patients of advanced maternal age or with recurrent miscarriage loss^16^.

However, problems can arise as some embryos may undergo more than one freeze-thaw cycle. The reasons for this include: (1) the clinical decision to thaw and subsequently perform PGT on an already cryopreserved embryo that has not undergone PGT before; or (2) embryos that underwent an initial round of PGT but failed to achieve a reliable result (e.g. amplification failure) requiring a subsequent round of PGT and vitrification^19^. There are several studies that have shown that multiple freeze-thaw cycles and biopsies negatively impacts pregnancy and live birth rate and leads to an increased rate of miscarriage^12–13, 17, 19–20^. These studies demonstrated that multiple freeze-thaw cycles and biopsies did not impact offspring health including offspring’s weight, crown rump length, or sex ratio outcomes^12, 17, 20^. However, a major limitation of these studies is the conflation of the effects of biopsy and cryopreservation, leaving the specific impact of vitrification alone on clinical outcomes unknown. This presents a knowledge gap and necessitates further research to isolate the effects of repeated cryopreservation and practices.

In light of this, the present study aims to investigate the isolated effects of repeated vitrification on embryo implantation potential and pregnancy rate. Specifically, we assessed the effects of multiple freeze-thaw cycles on the embryo, by examining cryosurvival rate and the impact on the two distinct cell lineages of blastocyst-stage embryos. Additionally, we investigated the impact of multiple freeze-thaw cycles on embryo implantation potential by transferring blastocyst-stage embryos into pseudo-pregnant female recipients and assessing pregnancy outcomes. Based on the literature and identified gap in knowledge, we hypothesised that repeated cryopreservation negatively impacts mouse embryo implantation potential and pregnancy outcomes. This study will contribute to a more refined understanding of embryo cryopreservation practices in ART, with the potential to inform and optimise infertility treatment strategies.

## Materials and methods

All reagents were purchased from Sigma Aldrich (St. Louis, MO, USA) unless stated otherwise.

### Animals and ethics

Female mice (CBA × C57BL/6 first filial (F1) generation; CBAF1, 21-23 days old), male mice (CBA × C57BL/6 F1, 6-8 weeks old) and female Swiss mice (6–8 weeks old) were acquired from Laboratory Animal Services (LAS) at the University of Adelaide South Australia, Australia. All mice were maintained in a controlled light-dark cycle (12 h light: 12 h dark cycle). Mice were fed with rodent chow and water *ad libitum*. All experiments were approved by the Animal Ethics Committee of the University of Adelaide (M-2022-101) and were conducted in accordance with the Australian Code of Conduct for Animal Care and Use for Scientific Purposes.

### Media preparation

The culture of cells was conducted using filtered media that were covered with a layer of paraffin oil (Merck Group, Darmstadt, Germany). Prior to use, the media were equilibrated for 4 h at 37°C in a humidified incubator with a gas mixture containing 5% O2, 6% CO_2_, and the remaining balance of N_2_. To ensure optimal conditions, all procedures were carried out on microscopes equipped with warming stages which were calibrated to maintain the temperature of the media in the culture dishes at a constant 37°C.

Sperm capacitation and in vitro fertilisation (IVF) procedures were performed in filtered Research Fertilisation medium supplemented with 4 mg/mL bovine serum albumin (BSA; MP Biomedical, Albumin NZ, Auckland, NZ). Resultant embryos were cultured in filtered Research Cleave medium (ART Lab Solutions, SA, Australia) supplemented with 4 mg/mL BSA.

For embryo vitrification and thawing, the base medium for embryo handling consisted of Research Wash medium (ART Lab Solutions, SA, Australia) supplemented with 5 mg/mL BSA (referred to as handling medium henceforth). The equilibration solution comprised of handling medium with 10% v/v ethylene glycol and 10%v/v dimethyl sulfoxide (DMSO). Vitrification solution was prepared by dissolving 1 M sucrose in the handling medium, supplemented with 16.6% v/v ethylene glycol and 16.6% v/v DMSO. Thawing solutions were prepared by diluting sucrose in the handling medium, with concentrations ranging from 0.3 M, 0.25 M, 0.15 M, to 0 M. After thawing, the embryos were allowed to recover in filtered Research Cleave medium supplemented with 4 mg/mL BSA for a duration of 6 h in the incubator at 37°C in a humidified incubator with a gas mixture containing 5% O2, 6% CO_2_, and the remaining balance of N_2_.

### Isolation of mouse gametes, in vitro fertilisation, and in vitro culture

Female mice were injected intraperitoneally (i.p.) with 5 IU of equine chorionic gonadotropin (eCG; Folligon, Pacific Vet Pty Ltd., Braeside, VIC, Australia), followed by and i.p. injection of 5 IU of human chorionic gonadotrophin (hCG; Pregnyl, Merck, Kilsyth, VIC, Australia) 48 h later.

Male mice were culled by cervical dislocation (13 h post-hCG) to collect the ductus deferens and the posterior segment of the epididymis in Research Wash medium. Spermatozoa were released from the vas deferens and epididymis (caudal region) by blunt dissection in 1 mL of Research Fertilisation medium, followed by capacitation (1 h), in a humidified incubator at 5% O_2_, 6% CO_2,_ 89% N_2_; 37°C.

At 14 h post-hCG, female mice were culled by cervical dislocation to collect the oviducts in pre-warmed Research Wash medium with 4 mg/mL BSA. A 29-gauge needle was used to puncture the ampulla to collect for expanded cumulus oocyte complex (COC).

Expanded COCs from each ampulla were then placed into pre-equilibrated dishes containing 90 µL drops of Research Fertilisation medium. Cumulus oocyte complexes were then co-cultured with 10 µL of capacitated sperm for 4 h inside a humidified incubator of 5% O_2_, 6% CO_2_ with a balance of N_2_ at 37°C. Four hours later, presumptive zygotes were cultured in Research Cleave medium with the density of 10 embryos per 20 µL drop of medium. Presumptive zygotes were incubated at 37°C in a humidified incubator of 5% O_2_, 6% CO_2_ with a balance of N_2_ and allowed to develop to the blastocyst-stage. Twenty-four hours post-fertilisation, the rate of development to the 2-cell stage was recorded, and blastocyst rate was recorded at 96 h post-IVF. Embryos at the two-cell stage were characterised by the presence of two blastomeres of equal size. Blastocysts were identified based on the presence of a blastocoel cavity that occupied at least two-thirds of the embryos size.

### Embryo vitrification and thawing

Embryo vitrification and thawing were performed as previously described^21^. Briefly, blastocyst-stage embryos (96 h post-IVF) were vitrified with the CryoLogic vitrification method (CVM). As shown in Figure 1, for each aim, blastocysts were separated into 4 experimental groups: fresh blastocysts (non-vitrified control), and blastocysts that underwent one, two or three freeze-thaw cycles. Following vitrification, embryos were thawed and recovered for 6 h to allow for re-expansion and revival in the incubator. Based on our prior experience and advice from clinical embryologists, a recovery time of six hours yields the highest survival rate for blastocyst-stage embryos. Cryosurvival rate was assessed following the 6 h recovery period for each freeze-thaw cycle. For each experimental group, embryos were separated into two cohorts to assess: (1) the number and viability of cells in the inner cell mass (ICM; fetal lineage) and trophectoderm (TE; placental lineage) of the blastocyst-stage embryo; and (2) post-embryo transfer outcomes (refer to **Fig. 1**).

**Figure 1.**
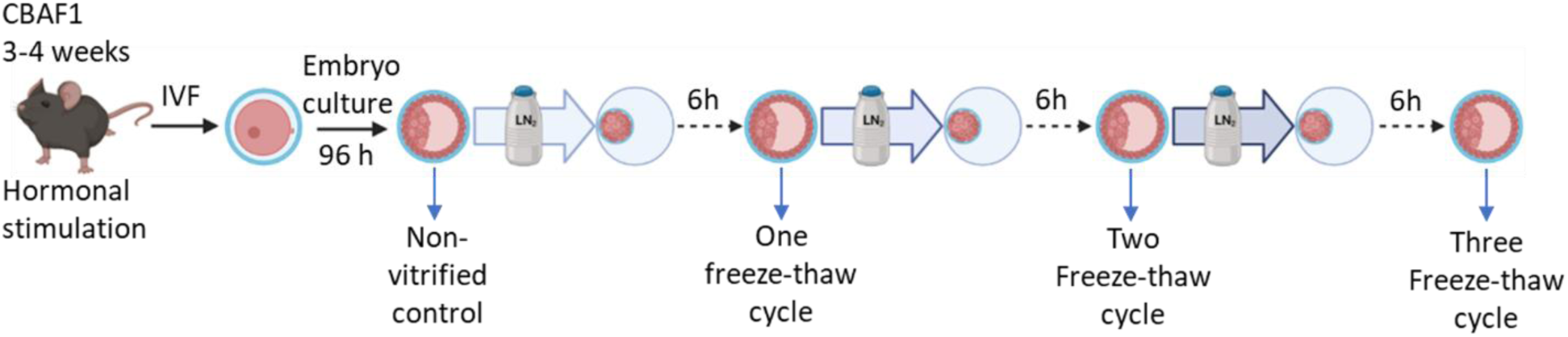
Illustration of experimental design. Briefly, blastocyst-stage embryos derived from IVF were separated into four experimental groups: (1) fresh (non-vitrified control); or subjected to (2) one freeze-thaw cycle; (3) two freeze-thaw cycle; (4) three freeze-thaw cycles. Embryos from each treatment group were then assessed for cryosurvival following 6 hours of recovery. Following scoring, embryos were assessed for the number of lineage-specific cells within the embryo and pregnancy outcomes following transfer. Image created using *Biorender*.

For vitrification, a four-well dish (ThermoFisher Scientific, Waltham, MA, USA) was filled with 600 µL of handling medium (HM), equilibration solution (ES), and vitrification solution (VS) (refer to **Fig. 2**). Media were pre-warmed to 37 °C for 15 min before vitrification. Embryos were rinsed twice in HM, then transferred into ES for 3 min. Embryos were then transferred into VS for 30 seconds and loaded onto a Fibreplug (CryoLogic, Pty. Ltd, VIC, Australia) with 2.5 µL of VS. Once loaded, the Fibreplug was immediately vitrified by touching the cryo block in liquid nitrogen and put into the Fibreplug straw. The Fibreplug straw was then stored within liquid nitrogen.

**Figure 2.**
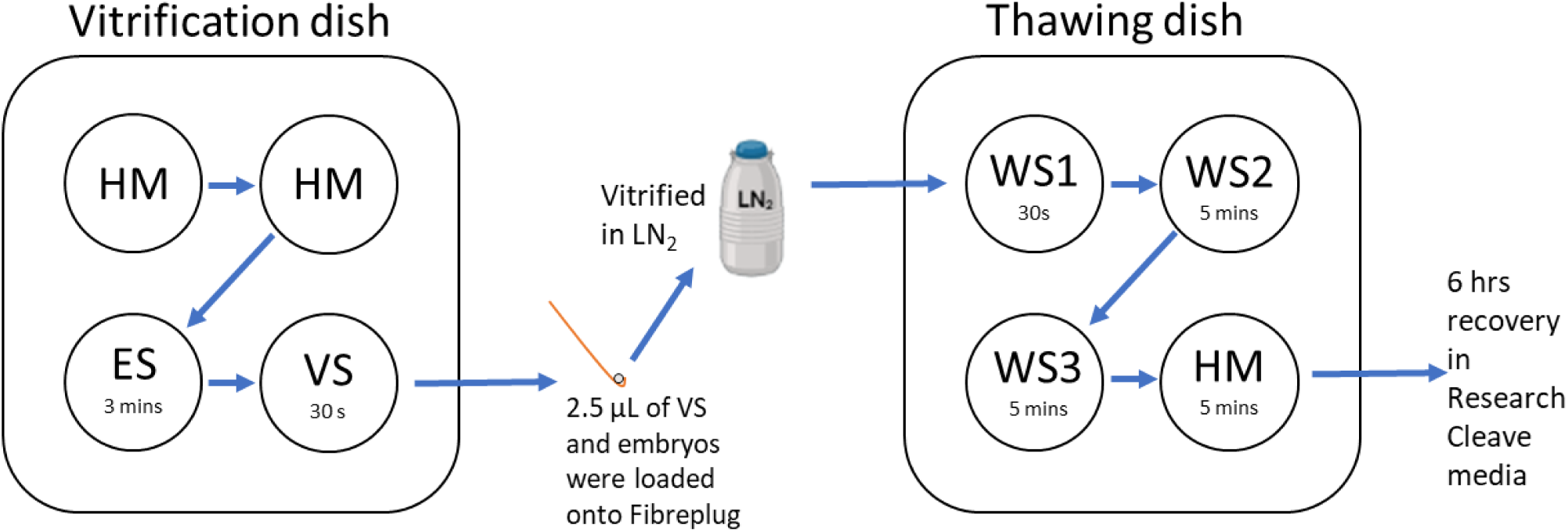
Schematic diagram for vitrification and thawing process. Vitrification dish contained: (1) Handling media (HM): Research Wash medium supplemented with 5 mg/mL BSA (referred to as handling medium henceforth); (2) Equilibration solution (ES): Handling medium with 10% v/v ethylene glycol and 10%v/v dimethyl sulfoxide (DMSO); and (3) Vitrification solution (VS): Handling medium with 1 M sucrose, 16.6% v/v ethylene glycol and 16.6% v/v DMSO. Thawing dish contained: (1) Warming media one (WS1): Handling medium with 0.3 M of sucrose; (2) Warming media two (WS2): Handling medium with 0.25 M of sucrose; (3) Warming media three (WS3): handling media with 0.15 M of sucrose; and (4) Warming media four (WS4): Handling media with no addition of sucrose. All vitrification and thawing processes occurred within the specified time as shown in the figure.

Thawing solutions were comprised of different concentrations of sucrose (0.3 M, 0.25 M, 0.15 M and 0 M) diluted in handling medium. For embryo thawing, a four-well dish was filled with 600 µL of handling medium supplemented with decreasing concentrations of sucrose (0.3M, 0.25M, 0.15 M and 0 M; referred as warming media 1, 2, 3 and 4, respectively). Media were pre-warmed to 37 °C for 15 min before thawing. Fiberplugs containing embryos were extracted from their straws and promptly immersed in a 0.3 M sucrose solution for a duration of 30 seconds, the embryos were then transferred sequentially into wells containing 0.25, 0.15 and 0 M sucrose for 5 min each. Finally, embryos were be transferred into Research Cleave medium and were allowed to recover for 6 h. Cryosurvival rate of thawed embryos was recorded at 6 h post-recovery (**Fig 2**).

### Embryo transfer

Prior to embryo transfer, vasectomy surgery was performed on male mice (CBA × C57BL/6 F1, 6-8 weeks old). Embryo transfers were performed as previously described^22^. Female Swiss mice were mated with vasectomised males, and the day where a copulation plug confirmed was recorded as day 0.5 post-coitum. Blastocyst-stage embryos following recovery were transferred into the uterine horns of pseudo-pregnant Swiss female mice at day 2.5 post-coitum. Sixteen morphologically normal, expanded blastocysts were transferred per mouse for 8 blastocysts allocated per uterine horn. Embryo transfers were performed on female Swiss mice under anaesthesia with 1.5% isoflurane maintained throughout surgery. Mice were monitored daily following embryo transfer surgery. Post-embryo transfer outcomes were recorded on day 18.5 post-coitum, which included pregnancy rate, viable fetal rate, resorption rate, fetal and placental weight.

### Immunohistostaining for fetal and placental cell lineage

Immunofluorescence for caudal-type homeobox 2 (CDX2) and octamer-binding transcription factor-3/4 (OCT-3/4) were used to assess the number of cells within the ICM and TE of a blastocyst-stage embryo, respectively. Embryos were fixed overnight in a 4-well dish with 400 µL of 4% paraformaldehyde (PFA; Emgrid, Australia), diluted in phosphate buffer saline (PBS). Embryos were then kept at 4°C in 0.3 mg/mL polyvinyl alcohol in PBS (PBV) until staining. All procedures for staining occurred at room temperature, unless stated otherwise. On the day of staining, embryos were first incubated with 0.1 M glycine for 5 min and washed with PBV prior to permeabilisation with 0.3% Triton X-100 for 20 min. Embryos were then washed three times with PBV and blocked with 400 µL of 3% BSA and 0.05% of Tween-20 in PBS (BSA_PSBT) for 1 h at room temperature. The embryos were then incubated with anti-OCT-3/4 mouse primary antibody (Santa Cruz Biotech, Dallas, TX, USA) at 1:100 dilution, and primary recombinant anti-CDX2 antibody (Abcam, Melbourne, AUS) at 1:500 dilution, both diluted in 10% goat serum at 4°C for 24 h. Following 24 h incubation, embryos were washed three times with PBV and incubated for 1 h in anti-mouse Alexa Fluor 488-conjugated secondary antibody (ThermoFisher, Waltham, MA, USA) and anti-rabbit Alexa Fluor® 594-conjugated secondary antibody (Life Technologies, Carlsbad, CA, USA). Both secondary antibodies were diluted at 1:500 dilution in 10% goat serum. Prior to imaging, embryos were washed with PBV and mounted in a 35mm glass bottomed dish within a 2 µL drop of PBV.

### Image acquisition and analysis

Images of OCT-3/4 and CDX2 immunostaining were captured on an Olympus FV10i confocal laser scanning microscope (Olympus, Tokyo, Japan) at 60× magnification using oil immersion compatible objective, with a numerical aperture equal to *NA* = 1.4. Images were acquired at regular intervals of 2 µm throughout the embryo, and a final z-stack projection was constructed. Samples were excited at a laser wavelength of 488 nm (emission detection wavelength: 490–525 nm) for OCT-3/4 and 594 nm (emission detection wavelength: 499–520 nm) for CDX2. The number of inner cell mass (ICM) cells and trophectoderm (TE) cells were quantified using OCT-3/4-stained and CDX2-stained cells, respectively. The percentage of ICM/TE was also calculated for each blastocyst.

### Statistical analyses

Statistical analyses were performed using GraphPad Prism Version 9 for Windows (GraphPad Holdings LLC, CA, USA) or Statistical Package for Social Science (SPSS) version 28.0.1.0 software. Data were subjected to normality tests prior to statistical analysis. Normally distributed data were analysed by an ordinary one-way analysis of variance (ANOVA) with Tukey’s multiple comparisons test. For data that did not follow a normal distribution, a Kruskal-Wallis test with Dunn’s multiple comparisons test was used for data analysis. Continuous data were presented as mean ± standard error of mean (SEM). Fetal weight and placental weight, and fetal: placental weight ratio were analysed by Linear Mixed model with litter size as covariant. Categorical data were expressed as percentages and analysed using the Fisher exact test. Specific details regarding the statistical tests employed can be found in the figure legends. Statistical significance was defined as a *P*< 0.05.

## Results

### Decreased cryosurvival and increased non-viable embryo rate with three freeze-thaw cycles

To determine the extent to which embryos retain their viability following repeated cryopreservation we assessed embryo viability at 6 h post-thawing. Cryosurvival was defined as successful re-expansion of the blastocoel cavity, while non-viable embryos were identified by fragmentation or collapse. There was a significant decrease in the cryosurvival rate and a significant increase in the number of non-viable embryos when embryos underwent three rounds of freeze-thaw cycles compared to those that underwent only one cycle (**Fig. 3 a and 3 b**; *P*<0.001). No significant difference was identified in cryosurvival rate or the rate of non-viable embryos for embryos that underwent two freeze-thaw cycles, compared to embryos that underwent one or three freeze-thaw cycles (**Fig. 3 a and 3 b**; *P* >0.05).

**Figure 3.**
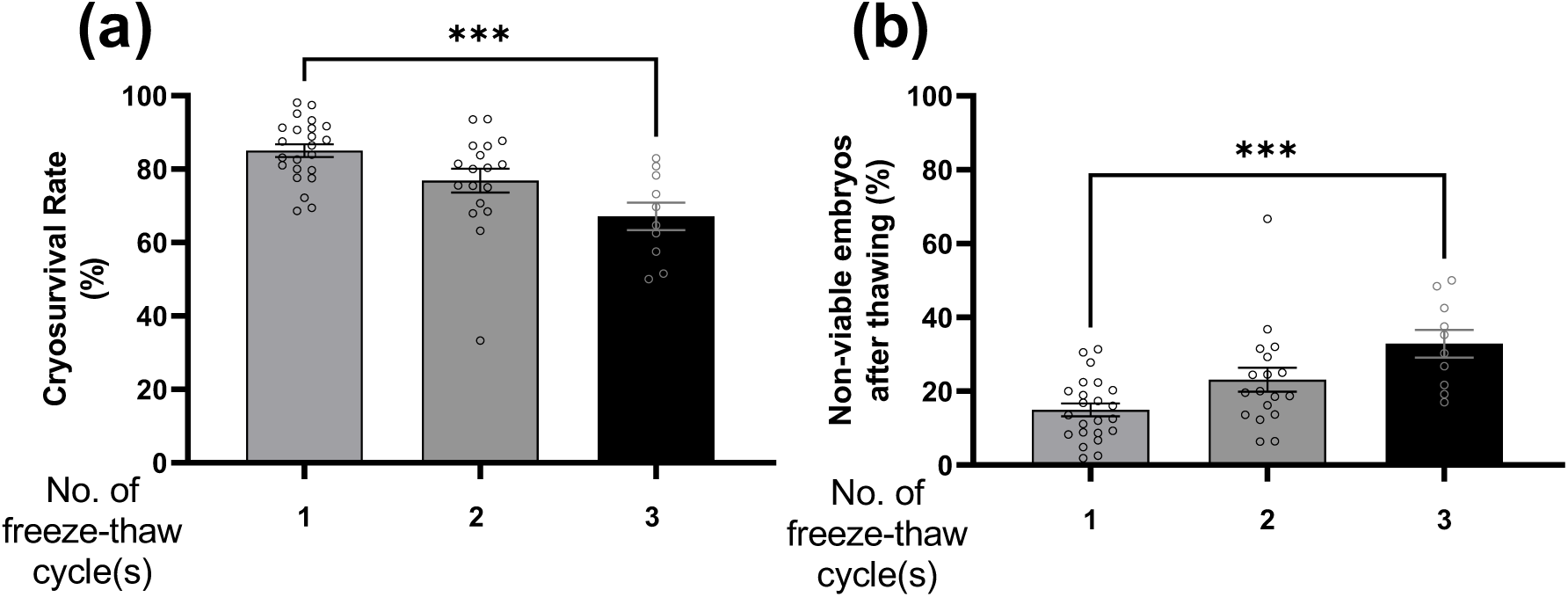
Increasing number of freeze-thaw cycles significantly reduced embryo cryosurvival and increased the number of non-viable embryos. Embryos were subjected to one, two or three freeze-thaw cycles. At 6 h following thawing, embryos were assessed by visual inspection and deemed to have survived (**a**: presence of an expanded blastocoel cavity) or were scored as non-viable (**b**: fragmented or collapsed blastocyst-stage embryos). Data are presented as mean ± SEM, n = 10-23 independent replicates; representative of 3035 (1 cycle), 1465 (2 cycles) and 447 (3 cycles) blastocysts per treatment group. Data were analysed using one-way ANOVA with Tukey’s multiple comparisons test. ***: *P*<0.001.

### Multiple freeze-thaw cycles reduced the number of cells in the inner cell mass in embryo

To evaluate the impact of multiple rounds of freeze-thaw cycles on the allocation of ucells to the two cell lineages within blastocyst-stage embryos we performed immunohistochemistry for the ICM (**Fig. 4 a, d, g, j**) and TE (**Fig. 4 b, e, h, k**). Following quantification, the number of ICM cells was significantly lower in embryos that underwent two and three rounds of freeze-thaw cycles compared to those that were not vitrified (**Fig, 4 m;** *P* <0.0001). However, there were no significant differences found between embryos that underwent one round of freeze-thaw cycle compared to those that were not vitrified (*P*= 0.0923). No statistically significant impacts on the TE cell number were identified among embryos that were cryopreserved or not cryopreserved (**Fig. 4 n;** *P* = 0.0661). However, the proportion of ICM over TE (ICM/TE, %) was significantly lower when embryos were cryopreserved compared to non-cryopreserved embryos (**Fig. 4 o;** *P* <0.05).

**Figure 4.**
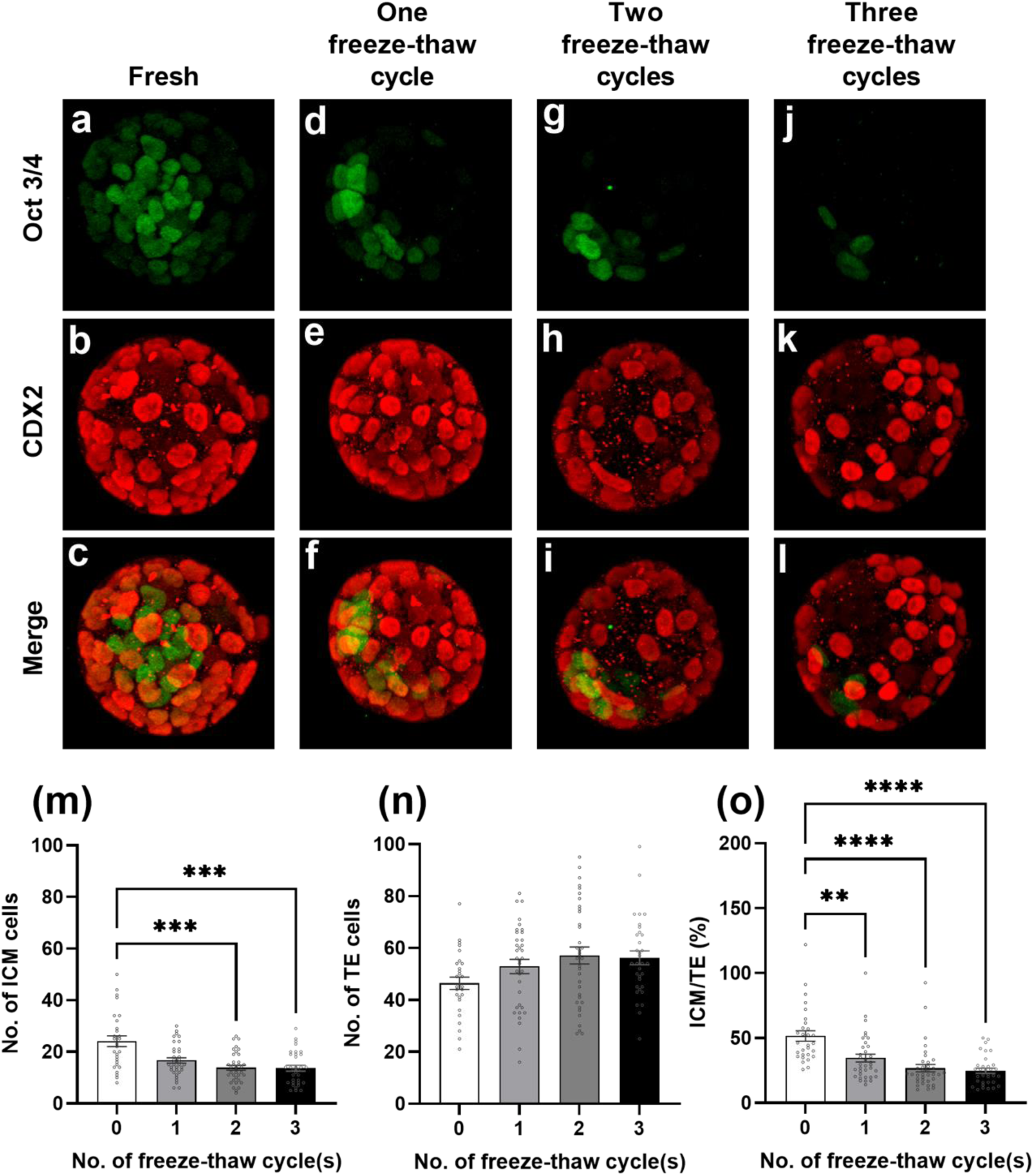
Increasing number of freeze-thaw cycles led to a significantly lower number of cells within the fetal lineage of the embryo. Blastocyst-stage embryos were either collected fresh (**a-c**), or subjected to one (**d-f**), two (**g-i**) or three freeze-thaw cycles (**j-l**). The number of cells within each lineage was assessed by immunostaining with Oct 3/4 (ICM; **a, d, g, j**) and CDX2 (TE; **b, e, h, k**). The number of cells within the ICM (**m**), TE (**n**), and the ratio of ICM/TE (**o;** expressed as a percentage) of resultant blastocysts for each treatment group were quantified. Data are presented as mean ± SEM, n= 28-36 embryos per treatment group, collected from 4 independent replicates. Data were analysed using Kruskal-Wallis test with Dunn’s multiple comparisons test. **: *P*<0.01, ***: *P*<0.001, ****: *P* <0.0001. ICM = inner cell mass, TE = trophectoderm.

### Performing multiple freeze-thaw cycles on the embryo does not impact pregnancy or viable fetal rates

A vital measure of the efficacy of cryopreservation techniques is their impact on pregnancy rates and fetal viability following transfer. To assess this, we transferred embryos that had undergone 0, 1, 2 or 3 freeze-thaw cycles into pseudo-pregnant female mice and assessed: (1) the pregnancy rate; (2) number of viable fetuses and (3) number of resorption sites. No statistically significant differences were found for pregnancy rate, number of implantations, viable fetuses or resorption sites (**Table 1**, *P* >0.05).

**Table 1.** Increasing numbers of embryo freeze-thaw cycles does not impact post-transfer outcomes. Results of embryo transfer experiments using blastocyst-stage embryos (96 h post-IVF) either collected fresh, or subjected to one, two or three freeze-thaw cycles. Embryo implantation and pregnancy outcomes for each treatment group was assessed at 18.5 d.p.c, by the percentage of viable and non-viable pregnancies, viable fetuses and resorption sites. d.p.c = days post-coitum. n = 12-17 individual mice per treatment group. Categorical data are described as percentages and compared using the Fisher’s exact test.

| Treatment | No. of females (%) |  |  | Viable fetal rate (%) |  |  |
| --- | --- | --- | --- | --- | --- | --- |
| No. of freeze-thaw cycles | Pseudopregnant Recipients | Pregnant (pregnancy rate; %) | Non-pregnant | No. of embryos implanted | No. of viable fetuses | No. of resorption sites |
|  | A | B (B/A) | C (C/A) | D | E (E/D) | F (F/D) |
| 0 | 22 | 12 (55) | 10 (45) | 64 | 29 (45) | 35 (55) |
| 1 | 29 | 13 (45) | 16 (55) | 70 | 38 (54) | 32 (46) |
| 2 | 33 | 14 (42) | 19 (58) | 60 | 18 (30) | 42 (70) |
| 3 | 43 | 17 (40) | 26 (60) | 53 | 18 (34) | 35 (66) |

### Fetal and placental weights not affected by the number of embryo freeze-thaw cycles

Considering that long-term health and developmental outcomes are paramount in assisted reproductive technologies, we extended our investigation to evaluate the impact of freeze-thaw cycles on fetal and placental weights. Our results showed that there was no impact on fetal weight or placental weight (**Fig. 5 a and 5 b**; *P* >0.05) with increasing numbers of freeze-thaw cycles. Additionally, we found that the number of times an embryo underwent a freeze-thaw cycle did not impact the fetal: placental weight ratio (**Fig. 5 c**; *P* >0.05).

**Figure 5.**
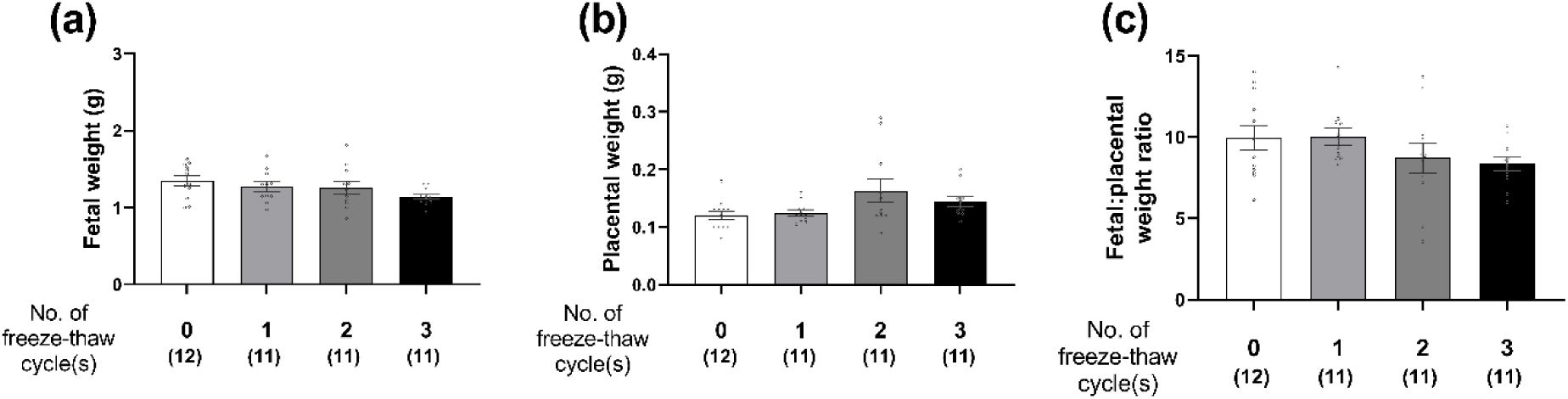
Increasing number of freeze-thaw cycles does not impact fetal or placental outcomes. Blastocyst-stage embryos were either collected fresh, or subjected to one, two or three freeze-thaw cycles and transferred into pseudo-pregnant mice. Fetal and placental outcomes were assessed on 18.5 d.p.c, by measuring fetal weight (**a**), placental weight (**b**), and calculated fetal: placental weight ratio (**c**). Data presented as mean ± SEM, n = 11-12 mice per treatment group: absolute numbers are shown in brackets. d.p.c = days post-coitum, analysed by Linear Mixed model with litter size as covariant.

### Discussion

Cryopreservation has emerged as an indispensable component of ART, particularly in conjunction with biopsy for PGT. However, when PGT fails, or in a case where a patient requires genetic testing on an already cryopreserved embryo, multiple rounds of biopsy and cryopreservation can occur. While previous studies have indicated that repeated biopsies and cryopreservation can adversely affect pregnancy and live birth rates^12–13^, the independent influence of multiple freeze-thaw cycles on clinical outcomes remains unknown, due to the confounder of concurrent biopsy^12–13^. Therefore, this study aimed to discern the singular effects of multiple freeze-thaw cycles on embryo health, implantation potential and resultant fetal health. We hypothesised that an increase in cryopreservation cycles would detrimentally impact the embryo health, implantation potential and fetal outcomes.

We found that increased numbers of freeze-thaw cycles is associated with a decline in cryosurvival and a higher number of non-viable embryos, indicating deteriorating embryo quality. Supporting this, recent literature in an ovine model has shown that vitrified embryos are more susceptible to cellular apoptosis and exhibit lower cryosurvival rates compared to their non-vitrified counterparts^14^. Furthermore, vitrification-induced cellular apoptosis of blastocyst-stage embryos is known to be associated with mitochondrial dysfunction, compromising the developmental potential of bovine and porcine embryos^23–24^. Given that apoptosis serves as a cellular survival mechanism to remove damaged cells^25^, and its association with embryo quality ^26^, our observations of increased non-viable embryos with increased freeze-thaw cycles, indicate that apoptosis may be occurring at a higher rate. However, it is beyond the scope for this project and will be the subject of a future study.

We observed a decrease in the number of cells within the ICM (fetal lineage) of embryos as the number of freeze-thaw cycles increased. This reduction may be attributed to an increased incidence of aneuploidy arising, which could then lead to apoptosis to eliminate these damaged cells. This is corroborated by a recent study demonstrating the active elimination of aneuploid cells in the ICM via apoptosis, while aneuploid cells in the TE (placental lineages) exhibit severe proliferation defects and are not removed^18^. This may explain our results on the number of TE cells (which were not impacted by increasing number of freeze-thaw cycles) where damaged cells in this cell lineage avoid apoptosis and are not removed due to aneuploidy. Whether the reduction in ICM cells in this study is mechanistically attributed to increased aneuploidy – via cryopreservation induced damage to the mitotic spindle of dividing cells – and then subsequent removal by apoptosis, is the subject of a future study. There is evidence to suggest that temperature fluctuations can cause microtubules in the oocyte to depolymerise, potentially leading to aberrant interactions between oocyte chromosomes, consequently increasing the incidence of aneuploidy in embryos post-fertilisation^27^. Given the rapid cellular divisions that preimplantation embryos undergo, it is possible that temperature variations during this critical phase may introduce errors, resulting in aneuploidy.

The proportion of ICM to TE cells is a recognised predictor of implantation and live birth, with a higher ICM to TE ratio associated with better pregnancy outcomes^28^. Consequently, the diminished ICM to TE ratio we observed in the embryos that underwent three freeze-thaw cycles could indicate decreased implantation potential. We hypothesise, that since the fetal lineage cells are critical for healthy fetal development, the embryo might be actively orchestrating the removal of damaged cells within the ICM to enhance developmental potential. Future studies could use genetic sequencing to study the ratio and integrity of damaged and apoptotic cells within ICM and TE and employ live/dead stains to study the distribution of apoptotic cells.

Remarkably, despite the deleterious effects on embryo quality, pregnancy rate, fetal and placental outcomes appeared to remain unaffected by increasing freeze-thaw cycles. This observation is in alignment with existing literature evaluating the combined effects of repeated cryopreservation with biopsy. These studies found embryos subjected to two freeze-thaw cycles and one biopsy exhibit lower implantation, pregnancy, and live birth rates coupled with an elevated miscarriage rates compared to embryos exposed to a single freeze-thaw cycle and biopsy^13, 17^. However fetal weight, height and sex ratio remained unaffected^20, 29^. Interestingly, although not significantly different, our results demonstrated that three freeze-thaw cycles had an approximately 42% lower rate of viable fetuses and 25% higher rate of resorption sites, when compared to non-vitrified control group. One plausible interpretation for our findings is the self-correction mechanisms in damaged embryos which involves^18, 30–31^: (1) autolysis of damaged embryos through miscarriage, irrespective of the proportion of compromised cells, allowing only less/non-damaged embryos to implant and develop into a fetus; or (2) increasing apoptosis rates to remove damaged cells, simultaneously increasing the proliferation of healthy cells to improve the implantation potential of embryo. Prior research indicates that post-implantation development can be rescued if sufficient euploid cells are present in the embryo^18^. We hypothesise that despite the reduction of cells in the ICM due to multiple freezing cycles, adequate numbers of viable cells may persist, facilitating the development of normal fetuses following the targeted removal of non-viable cells^18, 30–31^. Furthermore, given that embryos were allowed to recover for 6 h post-thawing in our study, these apoptotic and proliferative activities reached completion prior to embryo transfer, accounting for the lack of observed impact on pregnancy rates, or fetal and placental outcomes.

Moreover, the type of cryoprotectants used for vitrification may also impact embryo implantation potential. While sucrose, dimethyl sulfoxide (DMSO) and 1,2-propanediol (PROH), are ubiquitously used and have been reported to have no detrimental impact on the viability and morphology of oocytes and embryos^32–34^, there are contradictory studies that suggest that DMSO and PROH lead to premature centromere separation in murine oocytes and consequent embryo aneuploidy^35–38^. Furthermore, the concentration of cryoprotectants employed in clinical settings varies depending on the commercial vitrification kits used, such as: Vitrification Kit (Kitazato BioPharma, Shizuoka, Japan)^39^, Sage Vitrification Kit (Origio, Denmark)^40^ and Rapid-I devices (Vitrolife)^41^. Therefore, it may be plausible that the types and concentrations of cryoprotectant utilised in vitrification may be a modulating factor in clinical outcomes. Therefore, future investigations could systematically assess the influence of different cryoprotectant types and concentrations on IVF success. Such research could foster the optimisation of cryopreservation protocols, which is essential in assisted reproductive technology.

To the best of our knowledge, this is the first study to demonstrate the impact of multiple freeze-thaw cycles on embryo implantation potential and pregnancy rates. Future research could study how multiple freeze-thaw cycles affect the underlying survival mechanism in the embryo and explore avenues to ameliorate the adverse consequences of cryopreservation on embryo viability and development. The limitation of our study lies in the limited sample size for mice with successful implantation, therefore, future study should increase the sample size for pregnancy outcomes and improve the technique for embryo transfer. Moreover, it is crucial to recognise that this study was conducted in a murine model: therefore, data from this study should be interpreted with caution for different species. Further evaluation of the impact of repeated cryopreservation in preclinical studies and larger animal species are required.

In conclusion, our study underscores the critical insight that cryopreservation in isolation can influence embryo quality and calls for caution regarding the implementation of repeated cryopreservation cycles in IVF clinics. These findings hold clinical pertinence and have the potential to inform and shape future guidelines and practices concerning the application of embryo cryopreservation, ultimately contributing to the optimisation of outcomes in ART.

## Authors’ roles

KRD conceived the idea for the study. TCYT, RDR, and KRD were involved in experimental design. TL, RDR, TCYT and KRD were involved in interpretation of data. TL, SL and TCYT were involved in data acquisition, data analysis and generation of figures. TL, DJXC and TCYT wrote the first draft of most of the manuscript, RDR and KRD edited the final version of the manuscript. All authors critically revised and approved the final version of the manuscript.

